# Mutagenic impact and evolutionary influence of radiotherapy in hematologic malignancies

**DOI:** 10.1101/2024.11.15.623836

**Authors:** Benjamin Diamond, Dhanvantri Chahar, Michael D. Jain, Alexandra M. Poos, Michael Durante, Bachisio Ziccheddu, Marcella Kaddoura, Marios Papadimitriou, Kylee Maclachlan, Tomas Jelinek, Faith Davies, Nicholas B Figura, Gareth Morgan, Elias Mai, Katja C. Weisel, Roland Fenk, Marc S. Raab, Saad Usmani, Ola Landgren, Frederick L. Locke, Hartmut Goldschmidt, Jonathan H. Schatz, Niels Weinhold, Francesco Maura

**Affiliations:** Myeloma Institute, Sylvester Comprehensive Cancer Center, University of Miami, Miami, FL, USA; Lymphoma Service, Sylvester Comprehensive Cancer Center, Miami, FL, USA; Lymphoma Service, Moffitt Cancer Center, Tampa, FL, USA; Heidelberg Myeloma Center, Department of Internal Medicine V, Heidelberg University Hospital, Medical Faculty, Heidelberg University, Heidelberg, Germany; Clinical Cooperation Unit Molecular Hematology/Oncology, Department of Internal Medicine V, Heidelberg University Hospital, Heidelberg University, and German Cancer Research Center (DKFZ), Heidelberg, Germany; Myeloma Service, Memorial Sloan Kettering Cancer Center, New York, NY, USA; Department of Hemato-Oncology, University Hospital Ostrava and Faculty of Medicine, University of Ostrava, Czech Republic; Myeloma Research Program, NYU Langone, Perlmutter Cancer Center, New York, NY, USA; Department of Radiation Oncology, Moffitt Cancer Center, Tampa, FL, USA; Department of Oncology, Hematology and Blood and Marrow Transplant, University Medical Center Hamburg-Eppendorf, Hamburg, Germany; Department of Hematology, Oncology and Clinical Immunology, University-Hospital Duesseldorf, Duesseldorf, Germany

**Author notes:** **Corresponding Authors** Francesco Maura, MD, Myeloma Institute, Sylvester Comprehensive Cancer Center, University of Miami, 1120 NW 14th Street, Clinical Research Building, Miami, FL 33136; Niels Weinhold, PhD, Heidelberg Myeloma Center, Department of Internal Medicine V, Heidelberg University Hospital, Medical Faculty, Heidelberg University, Heidelberg, Germany.

**Keywords:** Whole Genome Sequencing, Mutational Signatures, Radiation, Radiotherapy, DNA damage, Multiple Myeloma, Diffuse Large B-cell Lymphoma

## Abstract

Ionizing radiotherapy (RT) is a widely used palliative and curative treatment strategy for malignancies. In solid tumors, RT-induced double strand breaks lead to the accumulation of indels, and their repair by non-homologous end-joining has been linked to the ID8 mutational signature in resistant cells. However, the extent of RT-induced DNA damage in hematologic malignancies and its impact on their evolution and interplay with commonly used chemotherapies has not yet been explored. Here, we interrogated 580 whole genome sequencing (WGS) from patients with large B-cell lymphoma, multiple myeloma, and myeloid neoplasms and identified ID8 only in relapsed disease. Yet, it was detected after exposure to both RT and mutagenic chemotherapy (i.e., platinum). Using WGS of single-cell colonies derived from treated lymphoma cells, we revealed a dose-response relationship between RT and platinum and ID8. Finally, using ID8 as a genomic barcode we demonstrate that a single RT-resistant cell may seed systemic relapse.

## INTRODUCTION

Ionizing radiotherapy (RT) is a clinical mainstay in the management of hematologic malignancies, in particular for local control and eradication of lymphoid malignancies. Depending on the specific disease entity and therapeutic intent, between 34 and 92% of patients with lymphoid malignancies will receive RT at some point in their disease course^1,2^. Ionizing RT results in the introduction of simultaneous DNA double-strand breaks (DSB) leading to cell death unless adequately repaired prior to cell division. Various endogenous repair processes, such as non-homologous end-joining (NHEJ), correct DNA damage but may leave mutagenic evidence of their activity within the genome that may be detectable as mutational signatures^3–11^. Recently, a higher indel (ID) burden has been observed in secondary tumors from patients exposed to RT and subsequent investigations have linked RT-induced DNA damage with the ID8 mutational signature^7,8^. Importantly, while RT is widely used in the clinical management of lymphoid tumors, its mutagenic impact on the mutational profiles and evolution of tumor cells at relapse has not been investigated. Furthermore, RT is known to increase the risk of clonal hematopoiesis (CH) and therapy-related myeloid neoplasms (t-MN), but it is largely unclear if this is driven by a direct mutagenic impact (i.e., accumulation of ID8 indels) or through other mechanisms^12^.

Therapy-related mutational signatures induced by select genotoxic chemotherapies are well-documented in post-treatment lymphoid and myeloid tumor samples^5,6,13–15^. As distinct mutagenic therapies have a predilection to cause single base substitutions (SBS) in preferred trinucleotide contexts and can be linked to a discrete clinical exposure, they have been used as powerful tools for reconstructing cancer evolution^6,13,16–19^. The measurement of therapy-induced mutations at the level of whole genome sequencing (WGS) is predicated by the expansion of a single cell that survives the exposure and sweeps its unique complement of therapy-induced mutations to clonal dominance. Leveraging this selective and mutagenic bottleneck, therapy-induced SBS signatures may assist in the timing of acquisition of driver events relative to therapeutic exposure in multiple cancer subtypes including therapy-related myeloid neoplasm (t-MN) and urothelial carcinoma^6,16,20^. With confidence in the source of ID8 in hematologic malignancies, it could similarly be used to track post-RT genomic evolution. ID8 has been observed in RT-related second-primary solid tumors^8^ and in somatic human cells^10^, animals, and cell lines following RT exposure^21,22^. However, ID8 has also been reported in patients not exposed to RT specifically with tumors with mutations in topoisomerase *TOP2A*^23^. The multi-etiological nature of ID8 therefore stands in contrast to the well-established chemotherapy-associated SBS signatures such as SBS31/SBS35 which are exclusively detected after exposure to platinum^5,7,9,15^. Overall, these data suggest that although RT leads to the acquisition of the ID8 indel signature, it is not the sole cause, and ID8 may more globally reflect errors introduced during repair of DSB by NHEJ in response to other genomic stresses^23^.

In this study, we leveraged a large series of 580 WGS to comprehensively assess the mutagenic impact of RT in multiple myeloma (MM), B-cell lymphoma (BCL), and t-MN. While the ID8 indel signature was expectedly enriched in samples collected following RT exposure, it was also detectable in cases exposed to chemotherapies including melphalan and/or platinum. To establish the relationship between ID8 and RT and ID8 and platinum-induced genomic damage, we treated a lymphoma cell line with cisplatin and RT and performed WGS on clonally expanded single cells. We demonstrate that exposure to genotoxic chemotherapies can produce a similar ID profile to RT that is indistinguishable in clinical samples. Using ID8 as a temporal barcode, we characterized relapse patterns of BCL and revealed that a single surviving cancer cell exposed to RT can seed a systemic relapse at an unrelated anatomic site.

## RESULTS

### Characteristics and Indel Profiles of Human Subjects

We interrogated the ID landscape of an assembled cohort of 580 WGS samples from 534 patients (**Fig. 1A**). Included in the total were 56 BCL tumor biopsies from patients treated with CD19 chimeric antigen receptor (CAR)-T cell therapy and a dataset of 100 BCL WGS from PCAWG^7,24^. Also included were bone marrow biopsies from 266 newly diagnosed MM (NDMM) from the GMMG-HD6 study and 98 relapsed/refractory MM (RRMM), including paired baseline/relapse samples from 61 patients^14,25,26^. We separately re-analyzed a set of t-MN (n=39) and 21 de novo AML to characterize t-MN transformation with regard to ID8^6,27^. All samples were aligned with genome reference GRCh37 and analyzed using paired germline samples. Median coverage for the relapsed LBCL, MM, and t-MN cases was 82x. Germline and tumor samples were collected under the respective Institutional Review Boards’ protocols and in accordance with the Declaration of Helsinki. Demographic data, disease features, and exposure history are listed in **Extended Data Table S1.** Detailed RT exposure history is reported in **Extended Data Tables S2-S4**. Of the 154 MM and LBCL samples collected at relapse, 107 (69.5%) had been exposed to melphalan, 53 (34.4%) had been exposed to platinum. 45 (29.2%) samples were from cases with prior RT exposure. Of these 40 (88.9%) had dual exposure to both mutagenic chemotherapy and RT while 5 samples (11.1%) were from patients with sole exposure to RT, without chemotherapy exposure.

**Figure 1.**
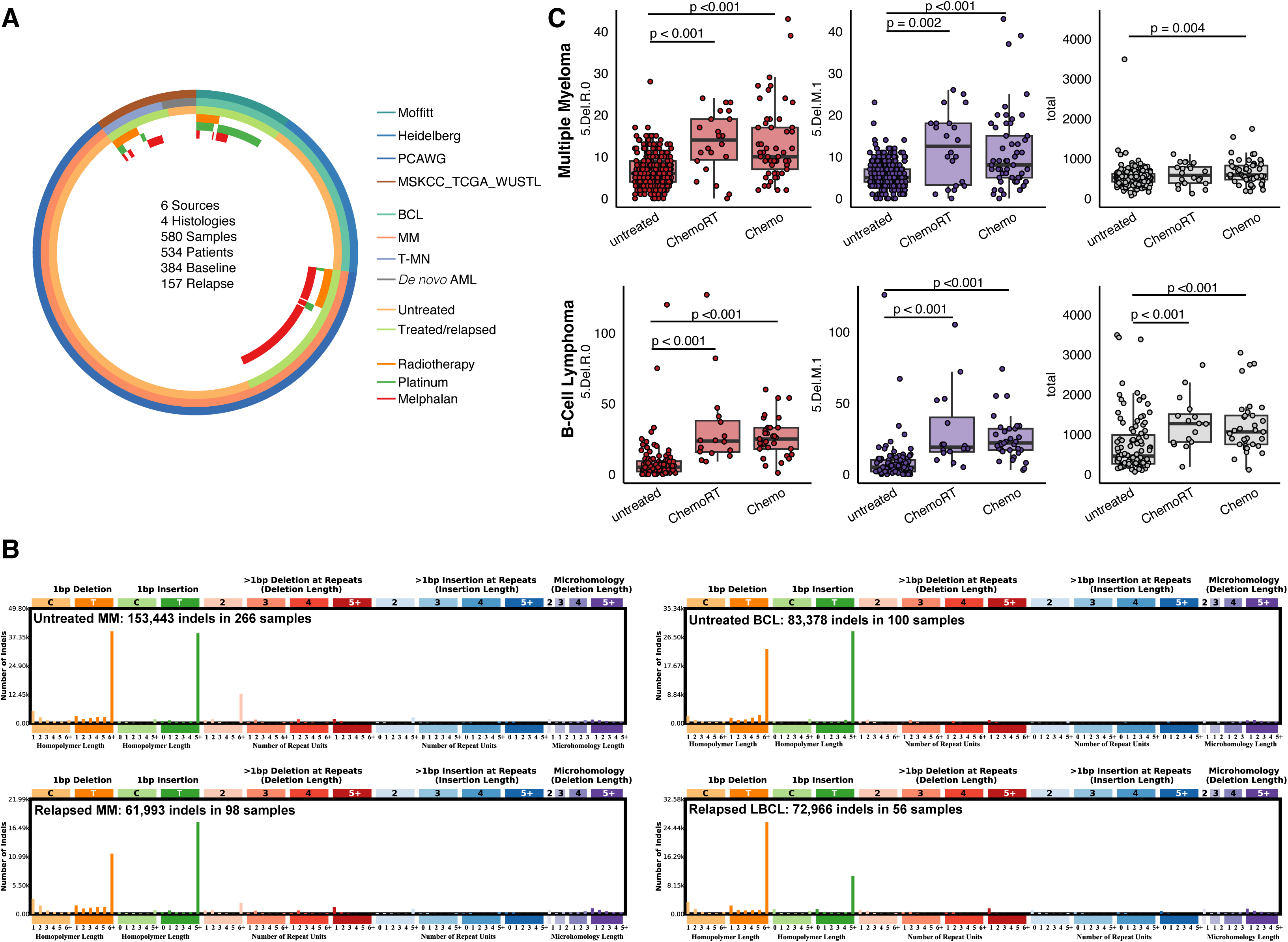
Indel Profiles of the Study Population. **A)** Summary of the WGS cohort. MSKCC; Memorial Sloan Kettering Cancer Center, PCAWG; Pan-Cancer Analysis of Whole Genomes, TCGA; The Cancer Genome Atlas, WUSTL; Washington University at St Louis. **B)** Cumulative ID profiles from MM and LBCL patients according to treatment phase with no gross difference in ID landscape. **C)** Differences in ID8/Radiotherapy-associated ID peaks (5.Del.R.0; 5+ BP Deletions at 0 Repeat Units, 5.Del.M.1; 5+ BP Deletions at Microhomology Length 1) and total ID counts between MM (top panel) and LBCL (bottom panel) according to radiotherapy and/or mutagenic chemotherapy exposure (i.e., platinum, melphalan). Two outliers were not plotted for graphical purposes (P334_BL, ID = 9,419; SP116697, ID = 11,193). *P* values were calculated using a Wilcoxon test. Box plot presents median ± 1^st^ and 3^rd^ quartiles.

We first defined the ID83 indel profiles of each patient (i.e., according to each of 83 indel subclasses)^7^. Without accounting for therapeutic exposure history, differences in the classes profiles across baseline and relapsed lymphoid cases were subtle (**Fig. 1B**). However, upon closer inspection, those samples that had been collected following exposure to either mutagenic chemotherapy and/or RT had a higher ID count at relapse in line with prior observations (**Fig. 1C, Supplemental Fig. 1A-B**). The observed increase in ID was concentrated in ID83 classes previously strongly associated with the ID8 indel signature (**Fig. 1C**)^4,7–10^. These are specifically large deletions (>5 base pairs) not restricted to sites of repeating units (5.Del.R.0) and also at regions of short microhomology length (5.Del.M.1). Both melphalan and platinum are known to induce DSB that, similarly to those induced by RT, may be repaired by NHEJ^28–32^. In line with this, the ID profiles of chemotherapy-treated tumors were similar to those of RT-treated tumors (**Supplemental Fig. 1A**).

### Enrichment of the ID8 Indel Signature in Chemoradiotherapy-exposed Lymphoid Malignancies

To define the ID signatures responsible for shaping the indel landscape of BCL and MM and the relationship between RT and ID8, we implemented a workflow consisting of *de novo* signature extraction, known signature assignment, and then refitting^33^. We first ran *SigProfiler* to extract and decompose both ID and SBS signatures agnostic to tumor histology^34^. Signature decomposition was further corroborated using a pairwise fitting contribution of COSMIC signatures to the *de novo* extracted signatures (https://github.com/UM-Myeloma-Genomics/Signature-Assignment)^6^. To confirm the presence of the de-convoluted signatures within our samples, we ran *mmsig* as a signature fitting algorithm, with modifications to accommodate both SBS96 and ID83 signatures (**Extended Data Table S5-6**)^15^. To maintain the highest stringency for ID8 calling, we accepted only ID8 fitting with non-zero bootstrapped confidence intervals estimated by *mmsig* (**Methods**)^35^. Overall, a total of 7 ID83 signatures were deconvoluted from 7 *de novo* extracted signatures (**Fig. 2**): ID1 and ID2; attributed to aging processes in all human cells^7^, ID4; attributed to RNAse H2 deficiency^36^, ID6; attributed to deficiencies in homologous recombination-based DNA-damage repair^7^, and ID8; attributable to the activity of NHEJ in response to DSB^28–32^. The latter two signatures, ID9 and ID12, are of unknown etiology.

**Figure 2.**
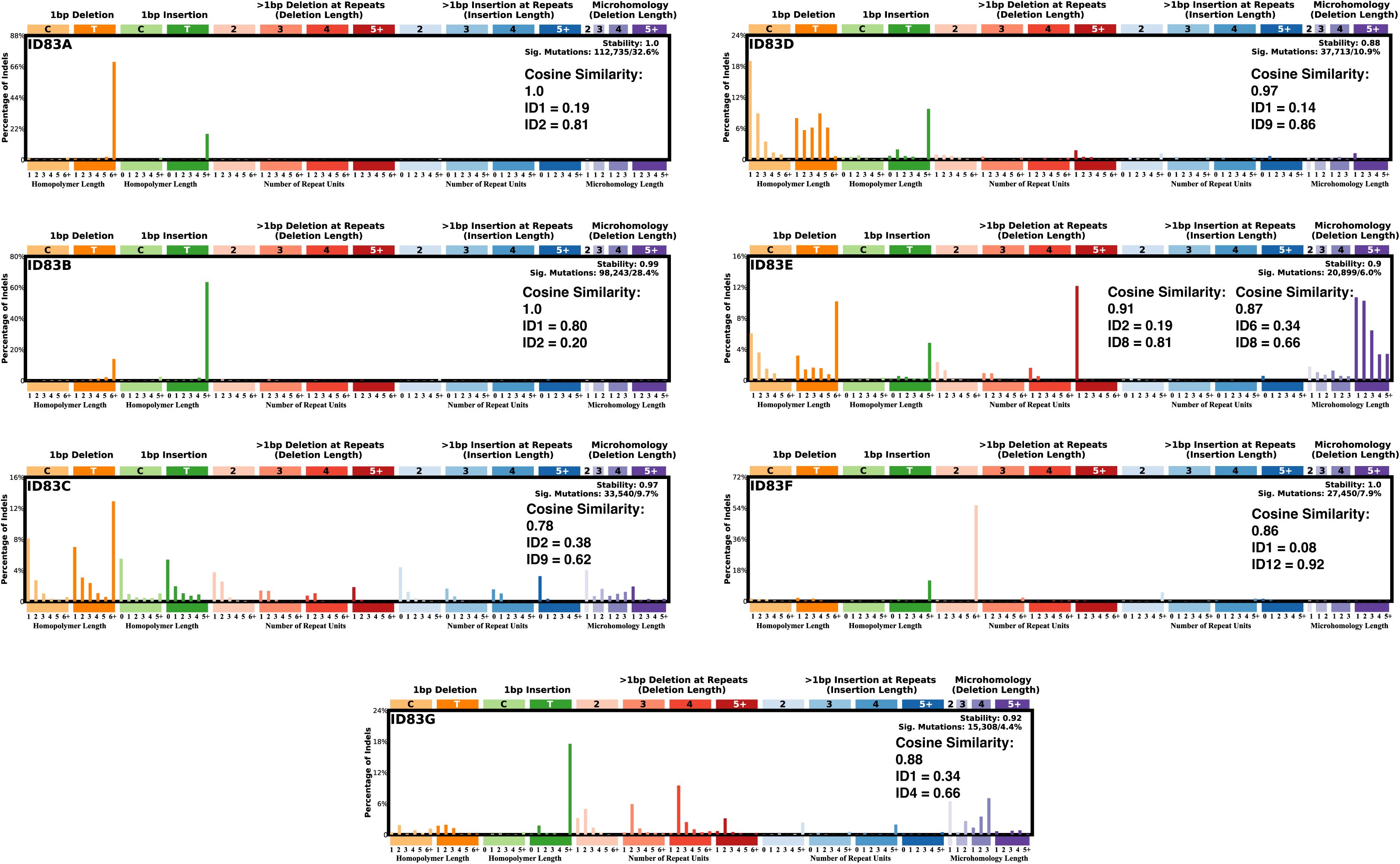
Indel Signatures Landscape of Lymphoid Malignancies. *De novo* extracted signatures for MM and LBCL cases using SigProfiler Extractor. Pairwise cosine similarities to known COSMIC signatures are denoted in the top right of each signature pane. The top two (e.g., highest cosine similarity) pairwise deconvolutions are included for ID83E.

While other indel signatures were represented regardless of treatment status (**Supplemental Fig 2A**-**B**), ID8 was seen only at relapse. Moreover, while the known associations of ID8 are with the repair of RT-induced DSB and rare p.K743N mutations in *TOP2A*, 10 of 18 (56%) ID8-positive samples were from radiation-unexposed patients without this *TOP2A* variant^23^. These cases all occurred in patients with prior mutagenic chemotherapy exposure (melphalan and platinum; **Fig 3A**). We therefore focused on the distribution of ID8 and its relationship with SBS chemo-related mutational signatures (i.e. platinum: SBS31, SBS35, E_SBS37; and melphalan:SBS99)^5,14,15^. Notably, ID8 was enriched in both MM and BCL samples with detectable SBS chemotherapy-associated signatures, regardless of RT exposure history (Wilcoxon rank-sum test, p<0.001; **Fig. 3B**). In fact, each ID8-positive case without prior RT exposure (8/18; 44%) had SBS evidence of chemotherapy-associated mutagenesis (**Fig. 3C**). On further inspection, even in those cases where ID8 was not called, chemotherapy signature-positive cases were seen to have an increased burden of the 5.Del.R.0 (Wilcoxon, p = 0.005) and 5.Del.M.1 peaks (Wilcoxon, p < 0.001) associated with NHEJ repair compared to those without chemotherapy-induced mutagenesis (**Supplementary** Fig. 3) suggesting that while mutagenic chemotherapy may induce ID8, its signal might be below limits of confident detection in presence of a low ID burden. Altogether, this suggests that the ID8 signature is therapy-related in lymphoid malignancies, and is limited to cases with prior RT exposure, and/or cases with evidence of chemotherapy-induced genotoxicity.

**Figure 3.**
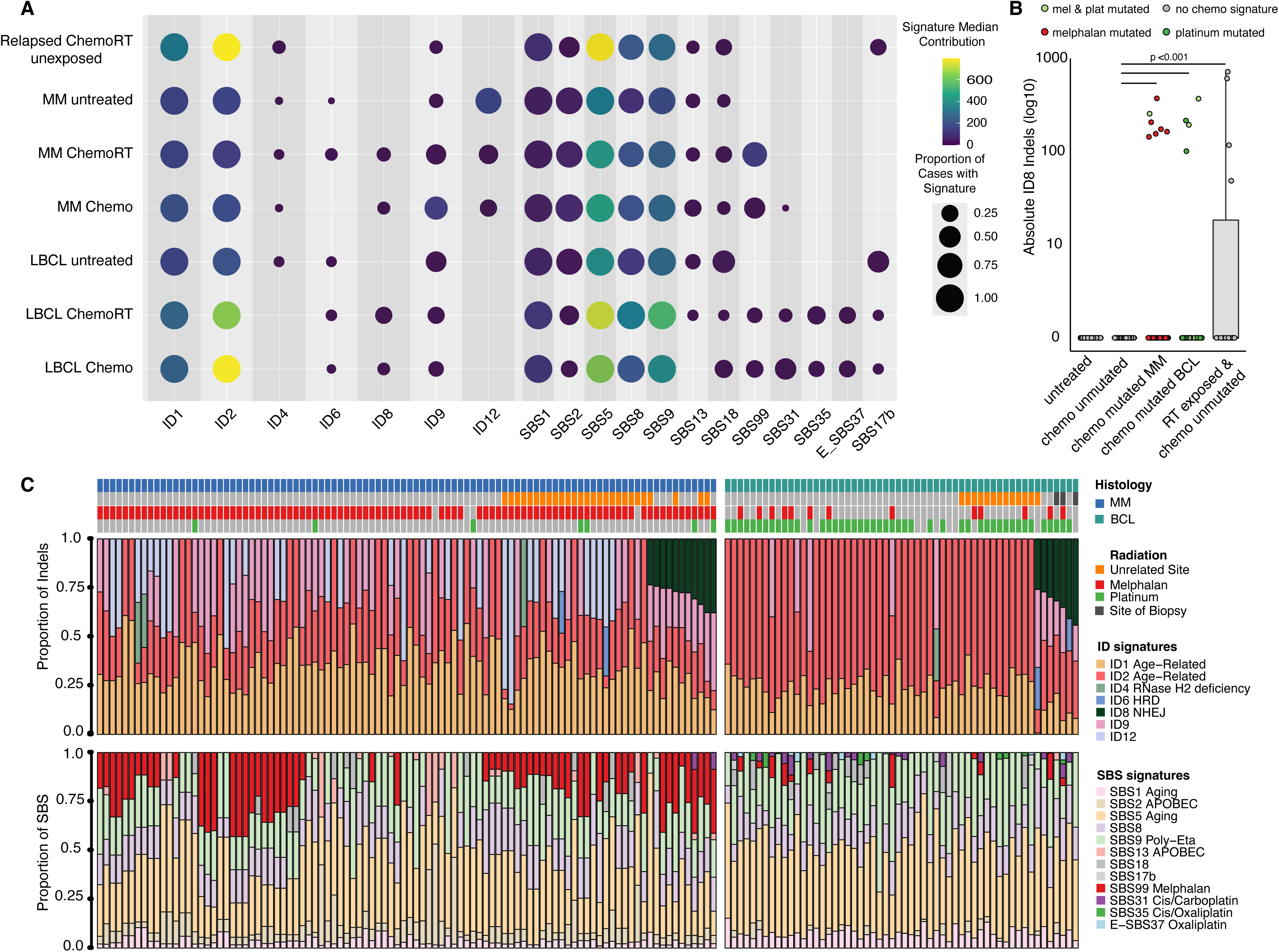
ID8 Occurs in RT-exposed and/or Chemotherapy-mutated Cases. **A)** Number of SBS and ID contributing to each lymphoid malignancy in the cohort. Tumors are grouped according to exposure to RT and mutagenic chemotherapy (i.e., platinum and melphalan). The size of each circle represents the number of cases positive for the signature and color corresponds to the median SBS/ID burden contributed by each signature. Relapsed_Unexposed are samples collected from relapse from both MM and LBCL without prior mutagenic chemotherapy exposure. **B)** ID8 indel counts among MM and BCL according to chemotherapy-associated mutation status. Chemo mutated cases have measurable SBS31, SBS35, E_SBS37, or SBS99 and may also have RT exposure. Chemo unmutated cases are relapse samples and may have been exposed to prior mutagenic therapy but do not bear the associated treatment-related signatures. *P* values were calculated using a Wilcoxon test. **C)** Mutational signatures contribution to the mutational burden of relapsed lymphoid malignancies. Top panels are the ID83 contribution and bottom panels are SBS96. Each column represents an individual samples. Samples are annotated by prior mutagenic chemotherapy exposure, RT exposure, and whether the sample collected for WGS was from a directly irradiated site of disease.

### Single-cell-derived colonies that survive platinum and RT exposure bear evidence of ID8

While this dataset suggests that DSB from mutagenic chemotherapy may be responsible for acquisition of ID8-signatures just as can be seen with RT, the contribution of either modality is unclear given that most patients have been exposed to concurrent or sequential chemotherapy and RT (88.9%; 40/45 samples). To evaluate the ability of RT and platinum to induce the ID8 signature, we exposed a DHL-4 *TP53*-mutant lymphoma cell line to daily doses of either of cisplatin or RT and selected individual resistant cells to find a new colony. For cisplatin experiments, DHL-4 cells were treated with a range of concentrations flanking the EC50 value (**Methods**). We chose DHL-4 as it is *TP53*-mutated and therefore more tolerant to DNA damage^37^.

After exposure to sequential doses of either therapy, viable cells were aliquoted and returned to the incubator for continued treatment, while a fraction was frozen to be used for downstream expansion (**Fig. 4A**, **Methods**). Mutagen exposure was continued until no viable cells were observed (Day Z). Proliferation was carefully monitored, and the maximum number of treatment days that allowed recovery of cell proliferation was defined as the sublethal dose (Day Z – 1). Half of this sublethal dose was defined as the medium dose. Frozen cells corresponding to these days were then thawed and single cells were then selected from medium- and sublethal RT and platinum populations and expanded to form single-cell derived colonies. As control, untreated cell line was subjected to the same single-cell expansion workflow (n=3 per group, total n=15, **Extended Data Tables S7-8**). We then performed bulk 60X WGS of the single-cell derived colonies (**Fig. 4A-4C**). Using the baseline untreated, unexpanded DHL-4 cells as the “germline” match, this strategy results in founding variants being filtered out as “normal” while allowing mutagen-induced variants, which are normally subclonal and unique to each exposed cell, to be fixed to clonality in a single-cell-derived colony (**Methods**)^9,38–40^. A feature of this approach is the overt selection for cells that maintain replication potential following multiple rounds of DNA damage. In fact, we noted that after just one to two doses of either of platinum or RT, cell division was halted, and required time for replication potential to be restored (**Fig. 4C**), potentially reflective of DNA-damage response and the loss of cells with excessive DNA damage. Overall, these dynamics likely reflect the biology of RT-treated cells in patients.

**Figure 4.**
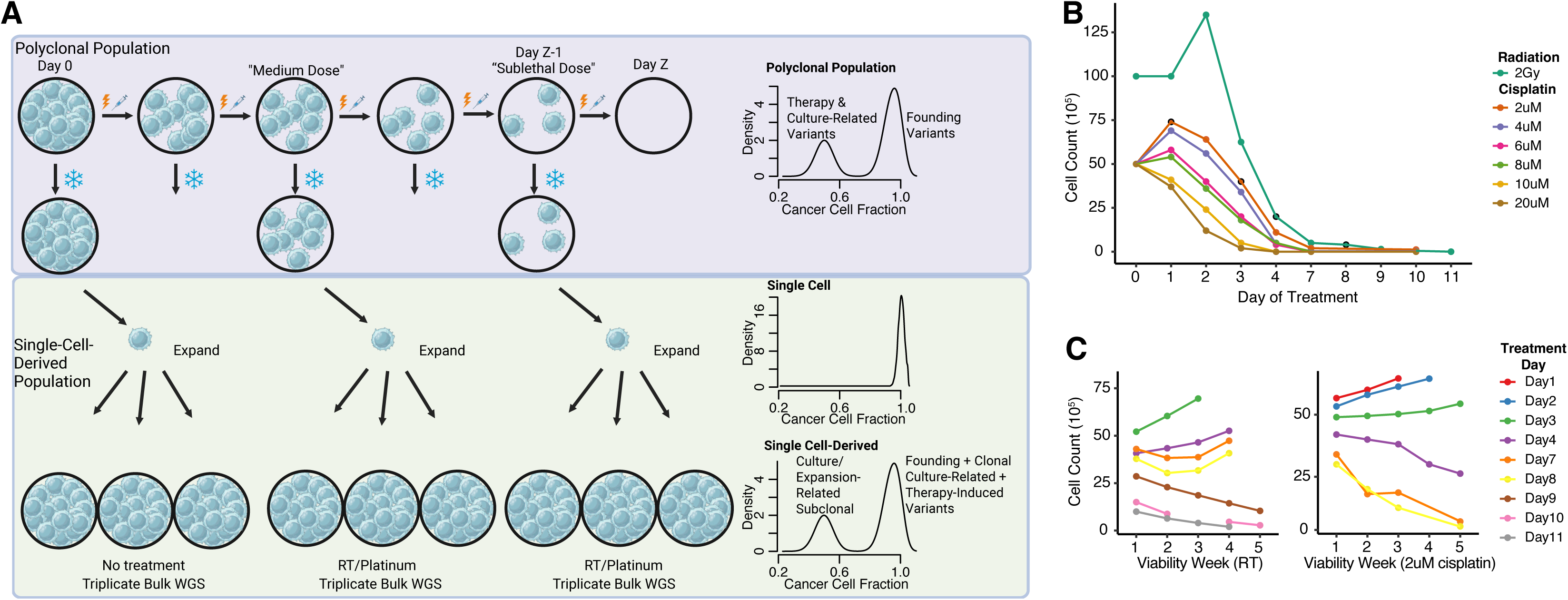
Clonal expansion of single cells exposed to platinum and radiotherapy. **A)** Cartoon summary for creation and bulk WGS of single-cell-derived colonies. The cumulative dose following which no recovery of cell division is observed is Day Z. *Created, in part, with bioRender.* **B)** Cell counts by exposure dose and day of treatment. Days chosen for single cell expansion are denoted with a black outline. **C)** Cells that survived daily doses of the highest concentration of cisplatin (2 µM) and still recovered replicative potential (Day 3) were designated as having undergone a sublethal dose, while the 2 µM treatment on Day 1 was classified as a medium dose. Cells exposed to RT that survived the maximum cumulative dose of 16 Gy and recovered replicative potential (Day 8) were classified as surviving the sublethal RT dose, with a cumulative dose of 8 Gy (Day 4) considered the medium RT dose.

We ran mutational signatures extraction, assignment, and fitting on clonal variants from 15 single-cell-expanded colonies. Four SBS96 (SBS96A-D) and three ID83 (ID83A-C) signatures were extracted *de novo* reflecting a composition of multiple established mutational signatures (**Fig. 5A**, **Supplementary** Fig. 4**, Extended Data Tables S9-10**). We next established a background mutational profile associated with single cell selection and expansion in experimental culture conditions using clonal variants from colonies derived from control untreated cells. The defined SBS-culture and ID-culture profiles reflect normalization of the SBS96 and ID83 variant profiles, respectively, from the three control colonies (**Methods**). Strikingly, these profiles were extremely similar to the extracted SBS96A (cosine similarity 0.978) and ID83A signatures (cosine similarity 0.987) suggesting they represent the background mutagenic impact of the experimental conditions (**Fig. 5A**, **Extended Data Table S10**). It was also apparent that SBS96B and ID83B were composed, in part, by the therapy-related signatures SBS31 (23% contribution) and ID8 (46% contribution) by pairwise fitting (**Methods**), respectively. Indeed, the number of SBS96B mutations increased in the platinum exposed colonies with evidence of a dose-response relationship from medium to sublethal doses of cisplatin (two sample t-test, p = 0.010, **Fig. 5B)**. Similarly ID83B indels increased from medium doses of RT to sublethal doses (two sample t-test, p = 0.037).

**Figure 5.**
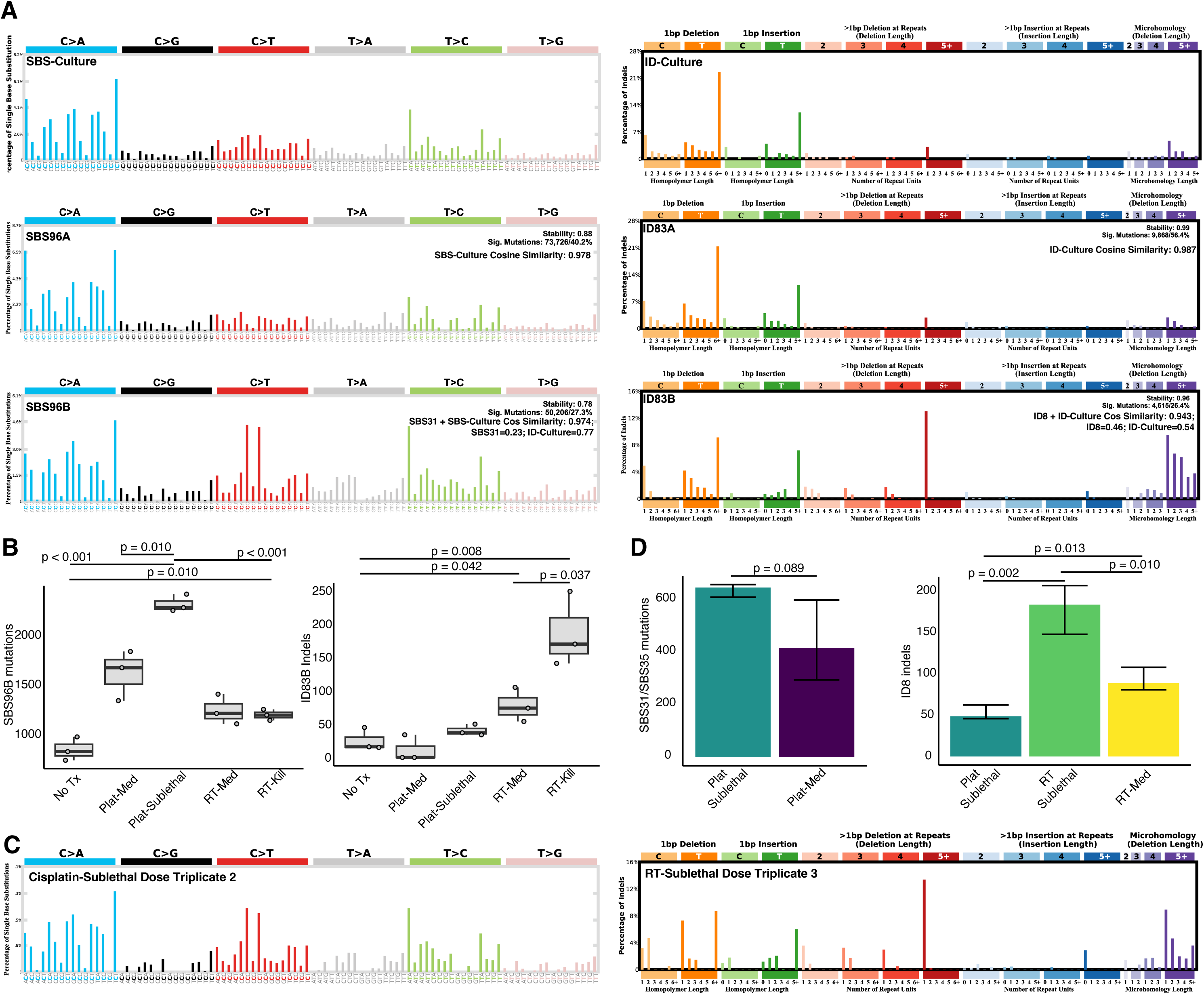
A) Selected de novo extracted signatures. SBS-Culture and ID-culture are the median SBS96 and ID83 profiles, respectively from the three control untreated single-cell-derived colonies. **B)** SBS96B (platinum) and ID83B (NHEJ) SBS and ID counts for single-cell-derived colonies demonstrating dose-response. P values reflect results of two-sample t-tests. Tx; Treatment. **C)** Selected de-noised SBS96 and ID83 profiles bearing similarity to SBS31/SBS35 and ID8, respectively. **D)** Therapy-related SBS31/SBS35 and ID8 mutation and ID counts from fitting of signatures to de-noised mutation and ID profiles.

We next performed a signal-to-noise (SNR) analysis for each of the treatment-exposed colonies as previously described by Kucab et al to provide clear evidence that the increases in SBS and ID burden are indeed due to cisplatin and RT and to facilitate the removal of culture-associated/background variants (**Extended Data Table S11**, **Methods**)^9,41,42^. These denoised profiles strongly resemble SBS31/SBS35 solely in cisplatin-exposed cases and ID8 in both cisplatin and RT-exposed cases (**Fig. 5C, Supplementary** Fig. 5-6). Moreover, when fitting ID8 to denoised profiles, a dose-response relationship was observed for both cisplatin- and RT-exposed colonies (**Fig. 5D, Extended Data Table S12**). Importantly, ID8 was detected in all three sublethal-dose platinum colonies; constituting strong validation that repair of platinum-induced DSB can yield the ID8 indel signature. However, ID8 was not fit in the medium-dose platinum colonies, suggesting that there is a minimum threshold of platinum exposure to yield enough DNA damage for a clear signature. The demonstration of a dose-response relationship provides a potential explanation for the relatively low penetrance of ID8 in melphalan and platinum-treated tumor samples, even among those with SBS-evidence of mutagenic chemotherapy.

Finally, we sought to determine whether the ID profiles induced by either of platinum or RT were distinguishable from one another. However, the median of denoised ID profiles from sublethal-dose platinum and RT colonies were similar and any differences in key peaks were not significant to differentiate the contribution of either source in samples with dual exposures (**Supplementary** Fig. 7).

### Impact of RT on therapy myeloid neoplasms development

Given the information gleaned here for lymphoid malignancies, we turned to a dataset of previously published t-MN where 19 of 39 patients had prior RT exposure for their primary malignancy (49%) to determine if similar patterns could be developed^6^. Running an ID signatures analysis with our updated workflow, we detected ID8 in only 2 t-MN (**Supplementary** Fig. 8**, Extended Data Tables S4 and S13**). One of these t-MN developed in a patient without RT, who was exposed to multiple rounds of platinum chemotherapy and PARP inhibition, while the other developed in a patient that was treated with RT and platinum-based therapy over 20 years prior for breast and ovarian cancers. Both of these tumors had evidence of platinum-associated mutagenesis (ie., SBS31/SBS35). As RT is a local therapy and t-MN arises from CH cells, the overall paucity of the ID8 signature in this cohort suggests that the mutagenic impact of RT on CH transformation may explain only a small fraction of t-MN cases. This implies that the increased t-MN risk following RT might be driven by other mechanisms that select CH for malignant progression.

### Tracking the progression of lymphoid malignancies in relation to therapeutic history

In lymphoid malignancies, RT may be used either for tumor palliation or with curative intent. In either case, however, the main goal is to provide local control and prevent local and/or systemic relapse. As ID8 can now reliably be associated with both genotoxic chemotherapy and RT, we sought next to reconstruct disease trajectory in relation to therapeutic exposures. We focused on three cases where ID8 could be clearly attributed to RT but not to other anti-cancer therapies (i.e. melphalan or platinum).

Patient CAR_285 was diagnosed with diffuse large BCL (DLBCL) and went on to receive RT to the brain and involved lymph nodes in the neck with achievement of local control. Five months later, however, the disease progressed at multiple systemic sites. A left posterior thigh mass was biopsied and subjected to WGS. Importantly the patient was platinum- and melphalan-naïve before tissue sampling. ID8 was present, indicating that ID8 indels were acquired as a consequence of RT and that a surviving RT-resistant cancer cell from the RT target regions seeded the mass in the thigh relapse (**Fig. 6A**)^5,43,44^.

**Figure 6.**
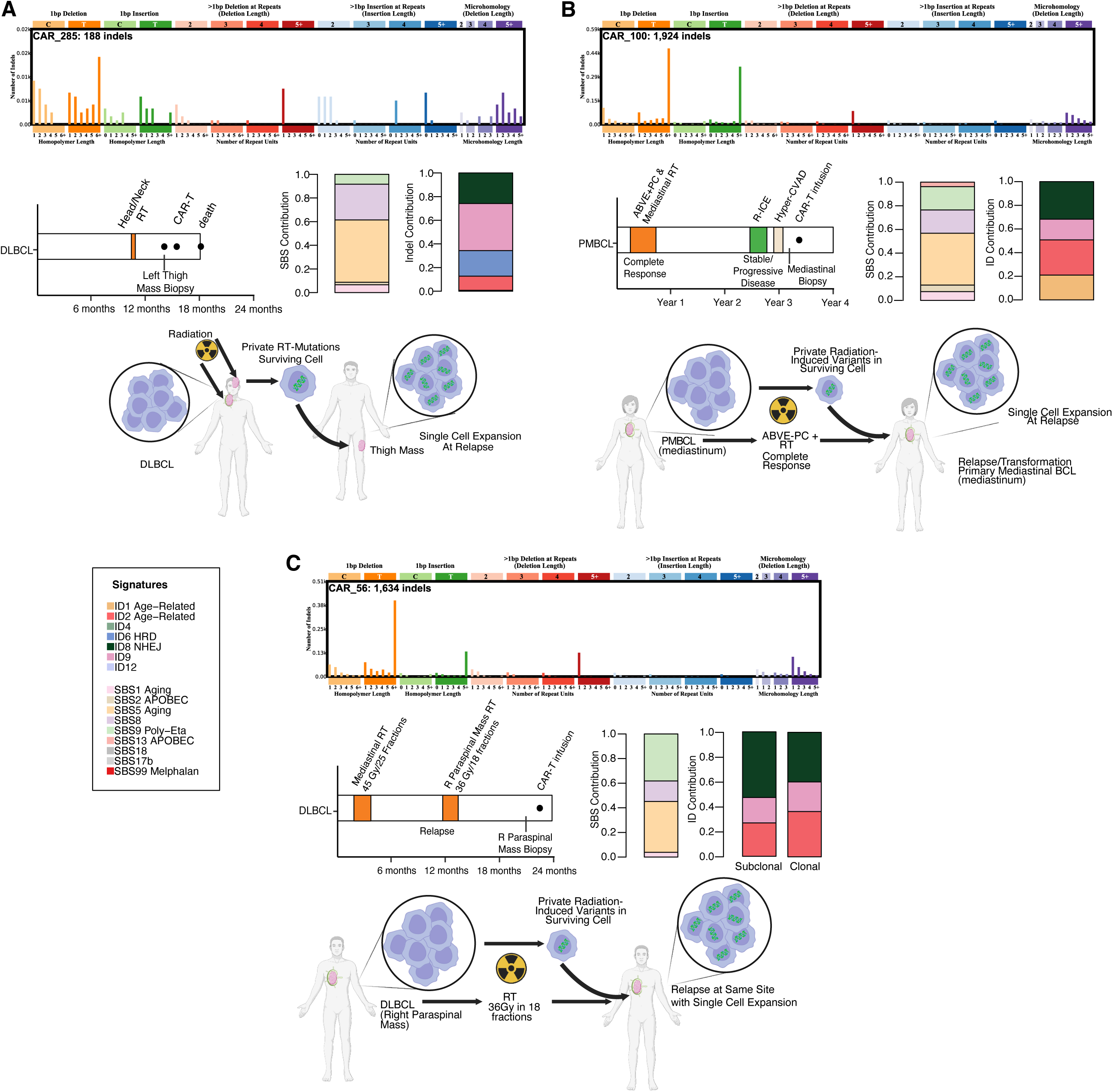
Vignettes demonstrating the use of the ID8 mutational signature to reconstruct lymphoid malignancy progression in the context of clinical exposure history for BCL (**A-C**). *Created, in part, with bioRender.* ABVE-PC; Doxorubicin Bleomycin Vincristine Etoposide Prednisone Cyclophosphamide, DLBCL; Diffuse Large B-Cell Lymphoma, Hyper-CVAD; cyclophosphamide vincristine doxorubicin dexamethasone methotrexate cytarabine, PMBCL; Primary Mediastinal B-Cell Lymphoma, R-ICE; Rituximab-Ifosfamide Carboplatin Etoposide;

Patient CAR_100 was originally diagnosed with primary mediastinal BCL and went on to receive combination chemotherapy with agents not known to increase mutational burden nor leave mutational signature evidence. Concomitant RT was delivered to the mediastinal mass with curative-intent. The combination of chemotherapy and RT (ChemoRT) resulted in a complete response but the disease relapsed at the same site 22 months later. As salvage, she underwent two cycles of platinum-based combination therapy that did not result in a tumor response. She then had further non-mutagenic combination therapy and then underwent re-biopsy of the mediastinal mass. WGS of the sample revealed ID8, consistent with an RT-resistant cell expanding to clonal dominance at local (i.e., involved site) relapse. Notably, there was no evidence of platinum-related signatures (i.e., SBS31/SBS35), most likely due to the lack of clinical response after platinum-based salvage therapies and the fact that these signatures are only detectable by bulk WGS in the case of single-cell expansions and subsequent subclonal selection (**Fig. 6B**)^5^.

Finally, Patient CAR_56 was diagnosed with DLBCL and subsequently received non-mutagenic chemotherapy and 45 Gy RT to the mediastinum in 25 fractions with curative intent. The disease relapsed systemically and 36Gy was delivered to a right paraspinal mass in 18 fractions. Another 10 months passed, and disease progressed further. Prior to CD19 CAR-T cell therapy, the formerly irradiated right paraspinal mass was biopsied. WGS of the sample revealed ID8, indicative either of a single cell seeding from the initial radiation exposure in the mediastinum or a local relapse of a surviving cell. However, the presence of the ID8 signature in both clonal and subclonal ID (**Methods**) can best be explained if a single cell from the initial mediastinal exposure seeded the paraspinal mass and contributed to the clonal ID8 indels. Subsequently, the second RT induced the acquisition of ID8 among different subclones (**Fig. 6C**).

## DISCUSSION

Despite impressive progress in recent years, most cancer patients remain to be treated with chemoRT-based regimens. In lymphoid malignancies, many LBCL will be exposed to combination chemotherapy consisting of mutagenic agents (e.g., melphalan or platinum) for consolidative or salvage approaches. Transplant-eligible patients with MM will be exposed to high-dose melphalan. In these contexts, RT is often used for control of symptoms (e.g. palliation in presence of spinal cord compression) or for eradication of residual disease. Despite the efficacy of these approaches, a significant fraction will experience a disease relapse and, in case of exposure to mutagenic agents, relapsed tumor cells usually bear the hallmarks of treatment-induced mutagenesis. The presence of chemotherapy-related SBS signatures is expected in most of these patients as the treatment is systemic, and mutagenic damage is often cell cycle independent^45^.

While SBS signatures are well characterized in the context of mutagenic chemotherapies, very little is known about how distinct anti-tumor agents affect the ID landscape at relapse in hematological cancers. Therefore, we assembled a large series of bulk WGS from both ND and RR lymphoid malignancies and we demonstrated that a specific ID mutational signature, ID8, is related to therapy-related genotoxicity from both RT and platinum and we leveraged it to gain resolution into patterns of relapse. To further validate this observation and to better understand the link between ID8 and platinum and RT, we used single cell-derived colonies from cell lines exposed to these mutagenic therapies. We confirmed the link between RT and ID8 and we revealed that platinum can also cause ID8 and produce ID profiles generally indistinguishable from the ones seen in RT-exposed cells. Interestingly, platinum- and melphalan-associated mutational signatures, when present, did not always co-occur with ID8. This can be explained either by limitations in the detectability of indel signatures (i.e., the number of indels is lower than SBS, reducing the accuracy of mutational signature extraction) or by other mechanisms related to the relationship between dose and differential DNA repair processes across tumors.

After unraveling the sources of ID8 in lymphoid tumors, a focus on patients without chemotherapy-associated mutagenesis allowed us to investigate genomic evolution and the dynamics of tumor relapse after RT. This is important as it is unknown if a relapse following RT is driven by a residual tumor cell outside of the RT field or by an exposed cell resistant to RT. Our data suggest that both of these scenarios occur. Importantly, we provide evidence that a single RT-resistant cell that survives treatment may be selected to seed systemic relapse at sites well outside the target field. These data are in line with observations from relapsed MM, where single cells can rapidly seed multiple organs, sometimes within just a few weeks^44,46^. Such patterns of relapse highlight the challenges in eradicating certain tumors, as a single surviving cell has the potential to promote tumor expansion and systemic progression.

Shifting the lens towards secondary malignancies that arise from prior chemoRT exposures, we observed that ID8 was rare in post-RT t-MN. Only 1 of 19 patients with prior RT exposure had evidence of ID8 in their t-MN, and ID8 may well have been introduced in this case by platinum chemotherapy. Although RT is associated with increased risk of CH and t-MN, our data suggest that the risk is not driven by a direct mutagenic role of RT but likely by other factors^12^. This aligns with observations of t-MN occurring after exposure to non-mutagenic agents, such as etoposide, suggesting that the origin of t-MN in the context of RT may lie in non-mutagenic mechanisms inclusive of therapy-induced selection of CH^6,27,47^. Future investigations may determine whether this is driven by the selection of fit clones, RT-induced inflammation and immunosuppression, and/or genetic predisposition.

Overall, the work presented here builds on the ability of therapy-related mutational signatures to understand disease-treatment interactions and reinforces the need for better strategies for tumor eradication, lest we preferentially select resistant disease to drive increasingly more aggressive relapses.

## METHODS

### Patients

Patient samples were collected from various publicly available sources. 39 t-MN were collected from EGAS00001006903, EGAD00001005028, and dbGaP (phs000159)^6,27^. 21 *de novo* AML came from dbGaP (phs000178). 256 ND MM from the GMMG-HD6 trial were taken from EGAS00001007469 and the remaining 10 NDMM and all RRMM were collected from EGAS00001006538, EGAS00001004363, EGAS00001004805 and EGAS00001005973. 100 LBCL from PCAWG are available at EGAS00001001692. The RR LBCL are available at dbGaP (phs003023.v1.p1).

### Patient sample processing and whole genome sequencing

Patient samples were processed as has been previously described^6,14,24,48,49^. However, a brief summary of processing of the contemporary datasets is provided here.

For the NDMM and RRMM from the GMMG-HD6 trial and Heidelberg University, mononuclear cells were collected from bone marrow aspirates (tumor) and from peripheral blood (normal) and were isolated using the Ficoll-Paque method. Bone marrow myeloma cells were enriched using human CD138 Microbeads and Manual MACS Cell Separation Units (Miltenyi Biotec, Bergisch, Gladbach, Germany). DNA of CD138-positive fractions was isolated using the Allprep Kit (Qiagen). WGS libraries were prepared with the Illumina TruSeq Nano DNA kit and sequenced on Illumina NovaSeq6000 Paired-end 150 bp S4. Raw sequencing data was processed and aligned to human reference genome build 37 version hs37d5 using the DKFZ OTP WGS pipeline^50^. ASCAT was used for copy number variants (CNV) and estimation of purity and ploidy. ID were called by Platypus^51^. SBS were determined using samtoolsmpileup (v1.2.166-3)^81^. For the ID and SBSs additional filtering steps were applied, including blacklist filtering^52^, fpfilter (https://github.com/genome/fpfilter-tool, only SBS) and removal of events located in regions coding for immunoglobulins. SV were called using SOPHIA (https://github.com/DKFZ-ODCF/SophiaWorkflow).

For the RR LBCL from Moffitt University, mononuclear cells from peripheral blood (normal match) or viably preserved tumor biopsies (tumor) had DNA extraction performed with AllPrep DNA/RNA Mini Kit (cat. no. 80284; Qiagen). Sequencing was performed with an Illumina NextSeq500. Sequencing libraries were prepared with the TruSeq DNA PCR-free HT sample preparation kit (Illumina). Libraries were normalized to 2.8 nM, and 24 samples were pooled for sequencing on a S4-300 flow cell on the NovaSeq 6000 and paired-end 150-bp reads were generated. Tumor and normal paired samples were aligned against the GRCh38 genome build and somatic single-nucleotide variants (SNVs) and short insertion-deletions variants (ID) were called using the DRAGEN Somatic Pipeline, Version 3.6.3. CNV were called with the Sequenza (version 3.0.0, https://github.com/cran/sequenza), as previously described for DLBCL. Manta was used to call SVs including deletions, inversion, translocations, and tandem duplications.

### Mutational signature analysis

Mutational signatures were defined and analyzed, in general, using a three-step process involving de novo extraction, assignment, and fitting^56,57^. *SigProfiler* (https://github.com/AlexandrovLab/SigProfilerExtractor) was utilized for the extraction of both SBS and ID signatures^58^. Assignment was performed first by cross-referencing the de-novo extracted signatures with the latest Catalogue of Somatic Mutations in Cancer (COSMIC) (https://cancer.sanger.ac.uk/cosmic/signatures/SBS) to identify the known mutational processes active within the cohort. For the ID signatures assignment, as previously described, we performed an adjusted deconvolution with the respective ID COSMIC catalogue using a bespoke algorithm (https://github.com/UMMyeloma-Genomics/Signature-Assignment)^33^. The code generates a pairwise fitting contribution of user-supplied reference mutational signatures to de novo extracted signatures and is particularly useful for the addition and evaluation of signatures not included in the COSMIC reference, and to minimize the assignment of rare and/or artifactual signatures. The top deconvolution combinations with biologic rationale reflective of signatures known to be active in included tumor histologies was chosen for each de novo signature. Subsequently, we employed *mmsig* (https://github.com/UM-Myeloma-Genomics/mmsig) as a fitting algorithm ^35^. This algorithm both confirms the presence of the assigned signatures using weights from the known clinical therapeutic exposures from a user supplied list and estimates the contribution of each mutational signature in every sample. Confidence intervals were generated by bootstrapping 1000 iterations. We modified the *mmsig* algorithm to accommodate ID83 signatures, as well as the previous SBS. Notably, this updated *mmsig* was used to re-analyze the ID83 signatures from the t-MN cohort.

Furthermore, for each patient sample, the cancer cell fraction (CCF) of each SBS and ID was calculated. On a sample-level, the median density of the CCF was used to split variants into clonal or subclonal. Mutational signatures were run on each of these subsets as above. At least 50 SBS were required for confident interpretation of SBS96 signatures and at least 40 ID for ID83 signatures.

### Statistical analysis and generation of plots

The Wilcoxon rank-sum test and Fisher’s exact test were used to compare the median of a continuous variable or the distribution of a discrete variable across groups, respectively. Two-sample t tests were employed to compare the difference in means between treatment groups. All p-values are two-sided, unless otherwise specified. Similarity between vectors was calculated by cosine similarity. All plots were generated in R-studio Version 4.1.0 (2021-05-18) in conjunction with Adobe Illustrator and Biorender.

### Creation of single-cell-derived colonies

Whole exome sequencing validated that our in-house DHL-4 cell line harbored a R273C *TP53* variant at 100% cancer cell fraction. To generate single-cell colonies post-radiotherapy, 10 million (0.5 million/mL) DHL-4 cells were irradiated with 2 Gy of cobalt-60 every 24 hours. Following each irradiation, cells were washed and re-cultured in fresh RPMI-1640 medium supplemented with 10% Fetal Bovine Serum (FBS), 5 μg/mL Plasmocin, 100 IU/mL Penicillin, and 100 μg/mL Streptomycin. Cells were maintained in an incubator at 37°C with 5% CO₂ throughout the experiment.

After each radiation exposure, approximately 50,000 viable cells were aliquoted and returned to the incubator. The cells collected 24 hours after the first radiation exposure were labeled as Day 1, and those collected after 48 hours, following the second round of exposure, were labeled as Day 2 and so on. Radiation exposure was continued until no viable cells were observed (Day Z). Proliferation was carefully monitored, and the maximum number of radiation cycles that allowed cell proliferation was defined as the sublethal dose (Day Z – 1). Half of this sublethal dose was defined as the medium dose.

For chemotherapy, cells were first treated with cisplatin to determine the IC50 (half-maximal inhibitory concentration). Based on the EC50 value of cisplatin in DHL-4 cells (9.02 µM) in viability assays, a range of drug concentrations flanking the IC50 value was selected for further treatment (2, 4, 6, 8, 10, 20 µM). Cells were treated every 24 hours with varying concentrations of either melphalan or cisplatin. After each 24-hour treatment, 50,000 cells were aliquoted and re-cultured in drug-free medium. Cells collected after the first 24-hour exposure were considered Day 1 dose, and those after 48 hours were considered Day 2 dose, continuing in this manner. The maximum treatment duration survived by the cells was labeled as the sublethal dose (Day Z-1), and half of this maximum duration was considered the medium dose.

For DNA extraction, the PureLink™ Genomic DNA Mini Kit (Invitrogen) was used following the manufacturer’s instructions.

### Single-cell-derived colony whole genome sequencing and post-processing

Sequencing was performed with an Illumina NovaSeqX to a target depth of 60X. All samples were first evaluated for concentration by Qubit (Thermo-Fisher) and for integrity by TapeStation (Agilent). Sequencing libraries were prepared with the TruSeq DNA PCR-free HT sample preparation kit (Illumina). The Covaris LE220 focus acoustic sonicator was used to fragment 1ug of total DNA to a target size of 350 bp. Size selection of blunt-end fragments was performed with AMPure bead purification (Beckman Coulter). A-base tailing was performed on the 3′ blunt ends, followed by adapter ligation and bead-based cleanup of libraries. Evaluation of library fragment size was done on the TapeStation (Agilent Technologies). Molarity quantification was performed by quantitative polymerase chain reaction with adapter-specific primers (Kapa Biosystems) on a Roche Light Cycler. Libraries were normalized to 2.8 nM, and samples were pooled for sequencing on a patterned flow cell on the NovaSeqX and paired-end 150-bp reads were generated. Tumor and normal paired samples were aligned against GRCh38 reference build and somatic single-nucleotide variants (SNVs) and short insertion-deletions variants (ID) were called using the DRAGEN Somatic Pipeline, Version 3.6.3. CNV were called with ASCAT. Manta was used to call SVs including deletions, inversion, translocations, and tandem duplications. For these analyses, the “normal” match was the base untreated and unexpanded DHL-4 cell line while “tumor” was each single-cell-derived colony (including untreated single-cell-derived controls).

### Signal-to-noise ratios and signature denoising

Based on work done by Kucab et al.^9^ we calculated a signal-to-noise ratio for each treated single-cell-derived colony. The SNR is calculated by finding the averaged Euclidean distance between the SBS or ID profiles of the treated colonies and the untreated single-cell-derived control colonies (SBS-culture; ID-culture) and then dividing this value by the variability (i.e., sum of the standard deviations) in the mutation or ID profiles. Based on prior work, an SNR>2 is considered to denote that the observed difference in mutation or ID profiles is not due to noise, but rather to the mutagen in question.

For each treated, single-cell expanded colony, as the SNR was larger than 2 in all cases, we subtracted the “noise (i.e., background mutations). Background mutations were characterized for each colony WGS by normalizing the extracted profiles of SBS96A and ID83A to the SBS or ID count, respectively, for each sample, and then taking the absolute value of the subtraction from each sample’s mutational/ID profile. SBS96A and ID83A were each nearly identical (cosine similarity>0.975) to the median mutational/ID profiles of the control (SBS-culture; ID-culture) which facilitated this denoising approach^41,42^. Mutational signatures analysis was then performed on denoised profiles as described above.

## DATA AVAILABILITY

All of the following data are publicly available:

- 39 t-MN were collected from EGAS00001006903, EGAD00001005028, and dbGaP (phs000159).
- 21 *de novo* AML came from dbGaP (phs000178).
- 256 ND MM from the GMMG-HD6 trial were taken from EGAS00001007469 and the remaining 10 NDMM and all RRMM were collected from EGAS00001006538, EGAS00001004363, EGAS00001004805 and EGAS00001005973.
- 100 BCL from PCAWG are available at EGAS00001001692. The RR LBCL are available at dbGaP (phs003023.v1.p1).

Upon publication of the manuscript, WGS data from the single-cell-derived colonies (n=15) will be available at EGA.

## CODE AVAILABILITY

All the code used for the WGS analyses has been uploaded to https://github.com/UM-Myeloma-Genomics.

## ACKNOWLEDGEMENTS

This work was supported by the Sylvester Comprehensive Cancer Center National Cancer Institute (NCI) Core Grant (P30 CA 240139). B.D. is supported by the American Cancer Society and is a K12 Scholar supported by the National Cancer Institute of the National Institutes of Health under Award Number K12CA226330. AMP is funded by the Medical Data Scientist Program of Heidelberg University, Faculty of Medicine.

The authors are grateful to the patients for participation in this study. The Heidelberg team thank the Sample Processing Lab, the High Throughput Sequencing unit of the Genomics & Proteomics Core Facility and the Omics IT and Data Management Core Facility of the German Cancer Research Center (DKFZ), the DKFZ-Heidelberg Center for Personalized Oncology (DKFZ-HIPO) office, the Biobank Multiple Myeloma UKHD and the Myeloma Registry for excellent services.

FM is supported by Leukemia and Lymphoma Society, Department of defense (DoD), and NIH-NCI.

## CONTRIBUTIONS

J.S., N.W., and F.M. conceived of, designed and supervised all experiments, performed the analysis, and wrote the paper. B.D., D.C., M.J., and A.P. analyzed the data, performed analyses, and wrote the paper. E.M, H.G., N.F., K.W., R.F., S.U., O.L, F.L., and M.R. contributed to patient consent and sample collection. M.D., M.K., M.P., B.Z., G.M., F.D., K.M., and T.J. analyzed the data and performed analyses.

## REFERENCES

1. Talamo G, Dimaio C, Abbi KK, et al. Current role of radiation therapy for multiple myeloma. Frontiers in oncology. 2015;5:40.

2. Thumallapally N, Meshref A, Mousa M, Terjanian T. Solitary plasmacytoma: population-based analysis of survival trends and effect of various treatment modalities in the USA. BMC cancer. 2017;17:1–11.

3. Chang HH, Pannunzio NR, Adachi N, Lieber MR. Non-homologous DNA end joining and alternative pathways to double-strand break repair. Nature reviews Molecular cell biology. 2017;18(8):495–506.

4. Kocakavuk E, Anderson KJ, Varn FS, et al. Radiotherapy is associated with a deletion signature that contributes to poor outcomes in patients with cancer. Nature genetics. 2021;53(7):1088–1096.

5. Pich O, Muiños F, Lolkema MP, Steeghs N, Gonzalez-Perez A, Lopez-Bigas N. The mutational footprints of cancer therapies. Nature genetics. 2019;51(12):1732–1740.

6. Diamond B, Ziccheddu B, Maclachlan K, et al. Tracking the evolution of therapy-related myeloid neoplasms using chemotherapy signatures. *Blood*, The Journal of the American Society of Hematology. 2023;141(19):2359–2371.

7. Alexandrov LB, Kim J, Haradhvala NJ, et al. The repertoire of mutational signatures in human cancer. Nature. 2020;578(7793):94-101.

8. Behjati S, Gundem G, Wedge DC, et al. Mutational signatures of ionizing radiation in second malignancies. Nature communications. 2016;7(1):1–8.

9. Kucab JE, Zou X, Morganella S, et al. A compendium of mutational signatures of environmental agents. Cell. 2019;177(4):821–836. e816.

10. Youk J, Kwon HW, Lim J, et al. Quantitative and qualitative mutational impact of ionizing radiation on normal cells. Cell Genomics. 2024;4(2).

11. Degasperi A, Zou X, Dias Amarante T, et al. Substitution mutational signatures in whole-genome–sequenced cancers in the UK population. Science. 2022;376(6591):abl9283.

12. Bolton KL, Ptashkin RN, Gao T, et al. Cancer therapy shapes the fitness landscape of clonal hematopoiesis. Nature genetics. 2020;52(11):1219–1226.

13. Bertrums EJ, Rosendahl Huber AK, de Kanter JK, et al. Elevated mutational age in blood of children treated for cancer contributes to therapy-related myeloid neoplasms. Cancer discovery. 2022;12(8):1860–1872.

14. Jain MD, Ziccheddu B, Coughlin CA, et al. Whole-genome sequencing reveals complex genomic features underlying anti-CD19 CAR T-cell treatment failures in lymphoma. *Blood*, The Journal of the American Society of Hematology. 2022;140(5):491–503.

15. Rustad EH, Yellapantula V, Leongamornlert D, et al. Timing the initiation of multiple myeloma. Nature communications. 2020;11(1):1917.

16. Sánchez-Guixé M, Muiños F, Pinheiro-Santin M, et al. Origins of second malignancies in children and mutational footprint of chemotherapy in normal tissues. Cancer Discovery. 2024;14(6):953–964.

17. Bertrums EJ, de Kanter JK, Derks LL, et al. Selective pressures of platinum compounds shape the evolution of therapy-related myeloid neoplasms. Nature Communications. 2024;15(1):6025.

18. de Kanter JK, Peci F, Bertrums E, et al. Antiviral treatment causes a unique mutational signature in cancers of transplantation recipients. Cell stem cell. 2021;28(10):1726–1739. e1726.

19. Van Hoeck A, Tjoonk NH, van Boxtel R, Cuppen E. Portrait of a cancer: mutational signature analyses for cancer diagnostics. BMC cancer. 2019;19:1–14.

20. Nguyen DD, Hooper WF, Liu W, et al. The interplay of mutagenesis and ecDNA shapes urothelial cancer evolution. Nature. 2024:1–10.

21. Adewoye AB, Lindsay SJ, Dubrova YE, Hurles ME. The genome-wide effects of ionizing radiation on mutation induction in the mammalian germline. Nature communications. 2015;6(1):6684.

22. Meier B, Volkova NV, Wang B, et al. C. elegans genome-wide analysis reveals DNA repair pathways that act cooperatively to preserve genome integrity upon ionizing radiation. Plos one. 2021;16(10):e0258269.

23. Boot A, Liu M, Stantial N, et al. Recurrent mutations in topoisomerase IIα cause a previously undescribed mutator phenotype in human cancers. Proceedings of the National Academy of Sciences. 2022;119(4):e2114024119.

24. Pan-cancer analysis of whole genomes. Nature. 2020;578(7793):82–93.

25. Mai EK, Goldschmid H, Miah K, et al. Elotuzumab, lenalidomide, bortezomib, dexamethasone, and autologous haematopoietic stem-cell transplantation for newly diagnosed multiple myeloma (GMMG-HD6): results from a randomised, phase 3 trial. The Lancet Haematology. 2024;11(2):e101–e113.

26. Cirrincione AM, Poos AM, Ziccheddu B, et al. The biological and clinical impact of deletions before and after large chromosomal gains in multiple myeloma. Blood. 2024;144(7):771–783.

27. Wong TN, Ramsingh G, Young AL, et al. Role of TP53 mutations in the origin and evolution of therapy-related acute myeloid leukaemia. Nature. 2015;518(7540):552-555.

28. Edler M, Jakubowski N, Linscheid M. Quantitative determination of melphalan DNA adducts using hplc—inductively coupled mass spectrometry. Journal of mass spectrometry. 2006;41(4):507–516.

29. Faivre S, Chan D, Salinas R, Woynarowska B, Woynarowski JM. DNA strand breaks and apoptosis induced by oxaliplatin in cancer cells. Biochemical pharmacology. 2003;66(2):225–237.

30. Sousa MM, Zub KA, Aas PA, et al. An inverse switch in DNA base excision and strand break repair contributes to melphalan resistance in multiple myeloma cells. PLoS One. 2013;8(2):e55493.

31. Ward J. The yield of DNA double-strand breaks produced intracellularly by ionizing radiation: a review. International journal of radiation biology. 1990;57(6):1141–1150.

32. Sears CR, Turchi JJ. Complex cisplatin-double strand break (DSB) lesions directly impair cellular non-homologous end-joining (NHEJ) independent of downstream damage response (DDR) pathways. Journal of Biological Chemistry. 2012;287(29):24263–24272.

33. Maura F, Degasperi A, Nadeu F, et al. A practical guide for mutational signature analysis in hematological malignancies. Nature communications. 2019;10(1):2969.

34. Islam SA, Díaz-Gay M, Wu Y, et al. Uncovering novel mutational signatures by de novo extraction with SigProfilerExtractor. Cell genomics. 2022;2(11).

35. Rustad EH, Nadeu F, Angelopoulos N, et al. mmsig: a fitting approach to accurately identify somatic mutational signatures in hematological malignancies. Communications biology. 2021;4(1):1–12.

36. Reijns MA, Parry DA, Williams TC, et al. Signatures of TOP1 transcription-associated mutagenesis in cancer and germline. Nature. 2022;602(7898):623–631.

37. Wang F, Liu J, Robbins D, et al. Mutant p53 exhibits trivial effects on mitochondrial functions which can be reactivated by ellipticine in lymphoma cells. Apoptosis. 2011;16:301–310.

38. Kuijk E, Jager M, van der Roest B, et al. The mutational impact of culturing human pluripotent and adult stem cells. Nature Communications. 2020;11(1):2493.

39. Lee-Six H, Øbro NF, Shepherd MS, et al. Population dynamics of normal human blood inferred from somatic mutations. Nature. 2018;561(7724):473-478.

40. Mitchell E, Spencer Chapman M, Williams N, et al. Clonal dynamics of haematopoiesis across the human lifespan. Nature. 2022;606(7913):343-350.

41. Huber AR, Pleguezuelos-Manzano C, Puschhof J, et al. Improved detection of colibactin-induced mutations by genotoxic E. coli in organoids and colorectal cancer. Cancer Cell. 2024;42(3):487–496. e486.

42. Pleguezuelos-Manzano C, Puschhof J, Rosendahl Huber A, et al. Mutational signature in colorectal cancer caused by genotoxic pks+ E. coli. Nature. 2020;580(7802):269–273.

43. Diamond BT, Ziccheddu B, Maclachlan KH, et al. Tracking the Evolution of Therapy-Related Myeloid Neoplasms Using Chemotherapy Signatures. Blood Journal. 2023:blood. 2022018244.

44. Landau HJ, Yellapantula V, Diamond BT, et al. Accelerated single cell seeding in relapsed multiple myeloma. Nature communications. 2020;11(1):3617.

45. Alexandrov LB, Nik-Zainal S, Siu HC, Leung SY, Stratton MR. A mutational signature in gastric cancer suggests therapeutic strategies. Nature communications. 2015;6(1):8683.

46. Rasche L, Schinke C, Maura F, et al. The spatio-temporal evolution of multiple myeloma from baseline to relapse-refractory states. Nature Communications. 2022;13(1):4517.

47. Pich O, Cortes-Bullich A, Muiños F, Pratcorona M, Gonzalez-Perez A, Lopez-Bigas N. The evolution of hematopoietic cells under cancer therapy. Nature communications. 2021;12(1):4803.

48. Maura F, Kaddoura MA, Poos A, et al. Temporal Genomic Dynamics Shape Clinical Trajectory in Multiple Myeloma. bioRxiv. 2024:2024.2008. 2030.610457.

49. Maura F, Rajanna AR, Ziccheddu B, et al. Genomic classification and individualized prognosis in multiple myeloma. Journal of Clinical Oncology. 2024;42(11):1229–1240.

50. Reisinger E, Genthner L, Kerssemakers J, et al. OTP: An automatized system for managing and processing NGS data. Journal of biotechnology. 2017;261:53–62.

51. Rimmer A, Phan H, Mathieson I, et al. Integrating mapping-, assembly-and haplotype-based approaches for calling variants in clinical sequencing applications. Nature genetics. 2014;46(8):912–918.

52. Danecek P, Bonfield JK, Liddle J, et al. Twelve years of SAMtools and BCFtools. Gigascience. 2021;10(2):giab008.

53. Martincorena I, Raine KM, Gerstung M, et al. Universal patterns of selection in cancer and somatic tissues. Cell. 2017;171(5):1029–1041. e1021.

54. Li Y, Roberts ND, Wala JA, et al. Patterns of somatic structural variation in human cancer genomes. Nature. 2020;578(7793):112-121.

55. Rustad EH, Yellapantula VD, Glodzik D, et al. Revealing the impact of structural variants in multiple myeloma. Blood cancer discovery. 2020;1(3):258–273.

56. Maura F, Degasperi A, Nadeu F, et al. A practical guide for mutational signature analysis in hematological malignancies. Nat Commun. 2019;10(1):2969.

57. Degasperi A, Amarante TD, Czarnecki J, et al. A practical framework and online tool for mutational signature analyses show intertissue variation and driver dependencies. Nature cancer. 2020;1(2):249–263.

58. Alexandrov LB, Kim J, Haradhvala NJ, et al. The repertoire of mutational signatures in human cancer. Nature. 2020;578(7793):94-101.

