## Supplemental Figures for "Mutagenic impact and evolutionary influence of radiotherapy in hematologic malignancies"

**Figure S1**: Mutational Profiles of Chemoradiotherapy-Treated Lymphoid Patients. **A)** Cumulative ID profiles of patients with relapsed disease having been treated with radiotherapy and/or mutagenic chemotherapy. **B)** Differences in SBS counts between MM (top panel) and LBCL (bottom panel) according to radiotherapy and/or mutagenic chemotherapy exposure (i.e., platinum, melphalan). Three outliers were not plotted for graphical purposes (SP116697, SBS = 112,529; P_308BL, SBS = 71,732; SP116668, SBS = 66901). *P* values were calculated using a Wilcoxon test. Box plot presents median ± 1st and 3rd quartiles.

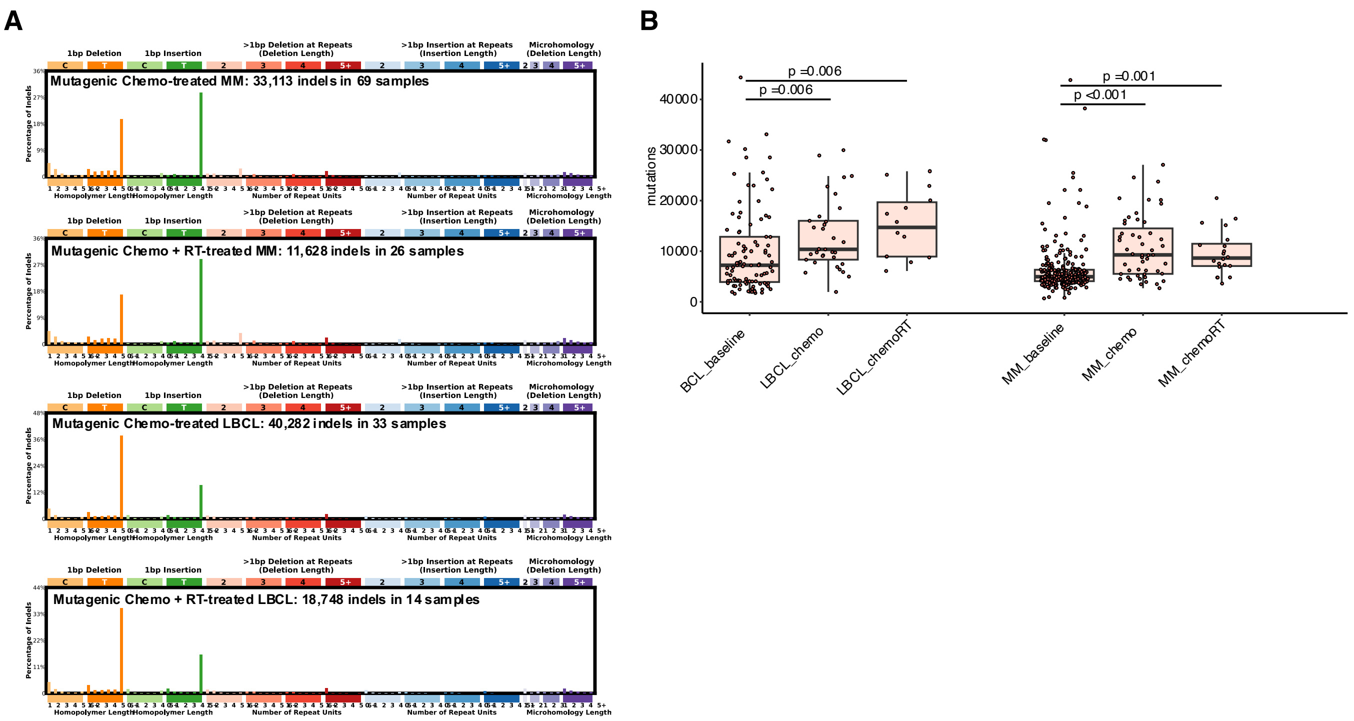

**Figure S2:** ID83 signature relationships to SBS96 signatures in newly diagnosed MM and BCL. **A)** Global SBS96 contribution for newly diagnosed MM. **B)** Indel signature landscape of untreated MM and BCL.

A

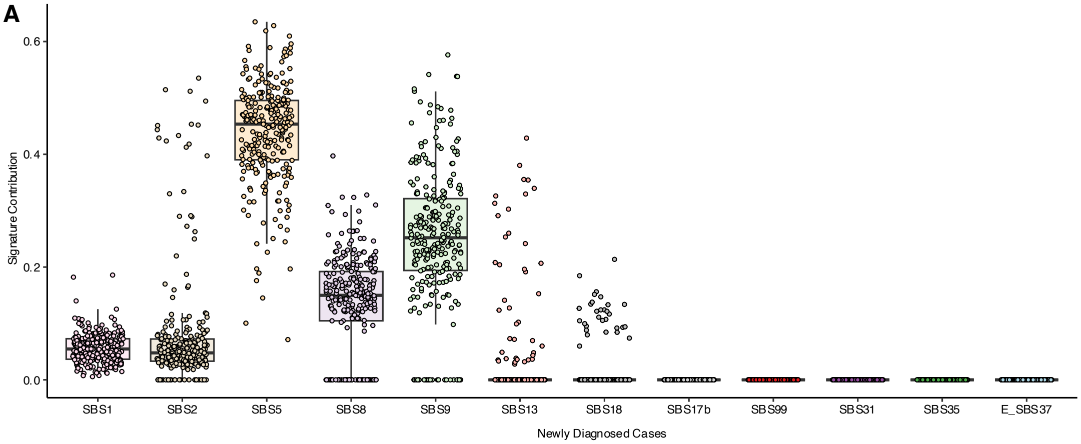


B

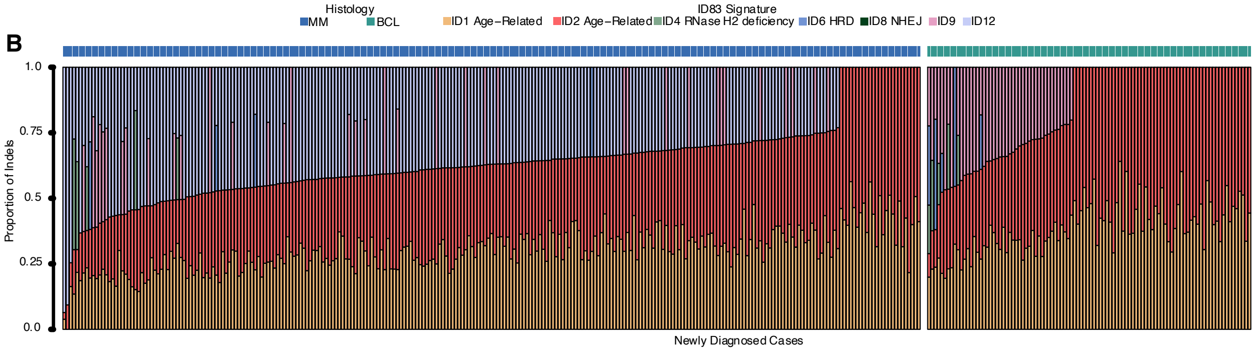

**Figure S3**: ID8-associated ID Peaks are enriched even in chemo mutated cases where ID8 was not called.

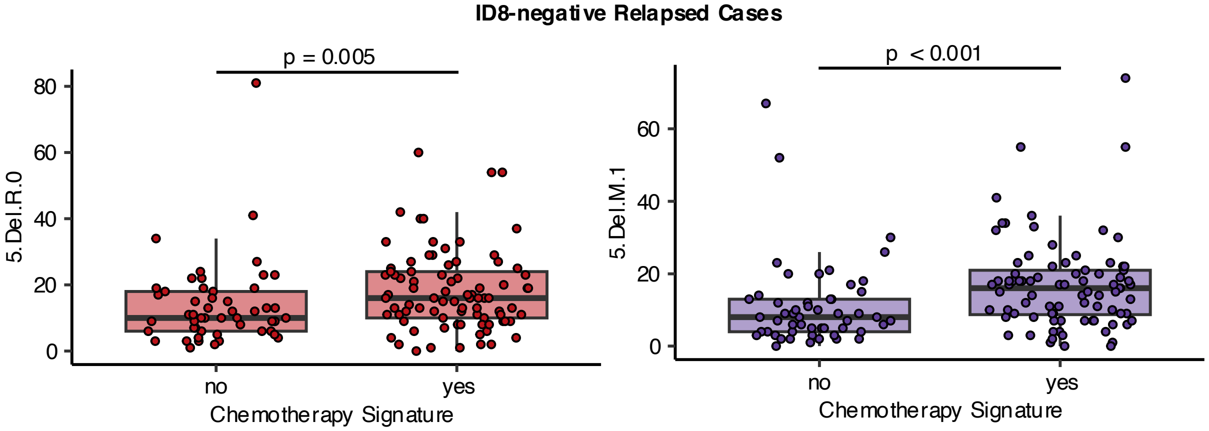

**Figure S4:** Remaining de novo signature extractions for single-cell-derived colonies not illustrated in Figure 4.

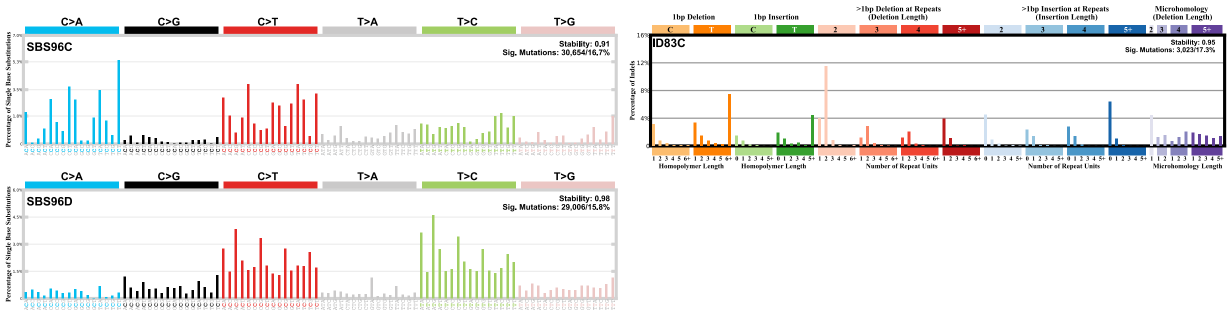

**Figure S5:** De-noised SBS96 profiles for single-cell-derived colonies.

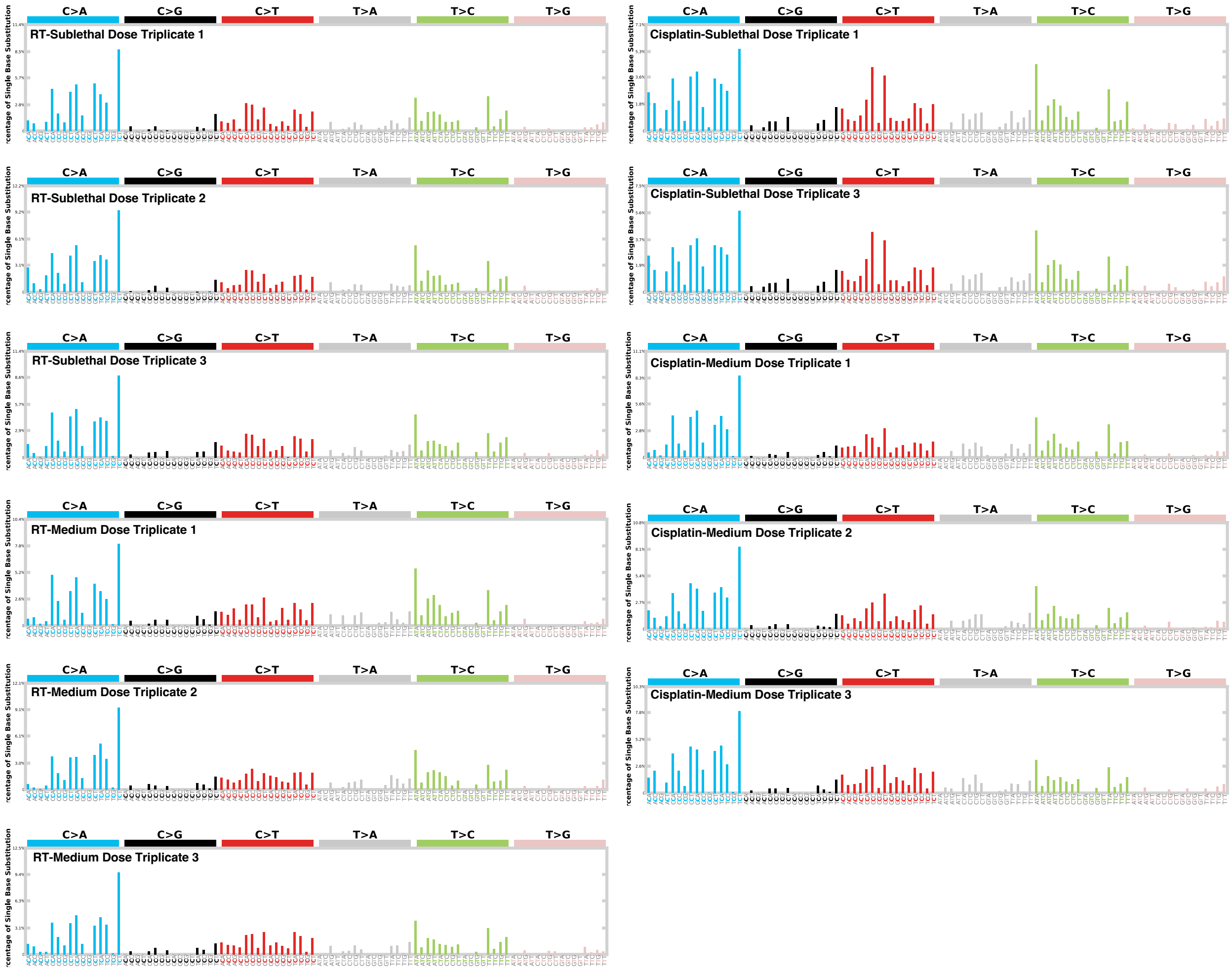

**Figure S6:** De-noised ID83 profiles for single-cell-derived colonies.

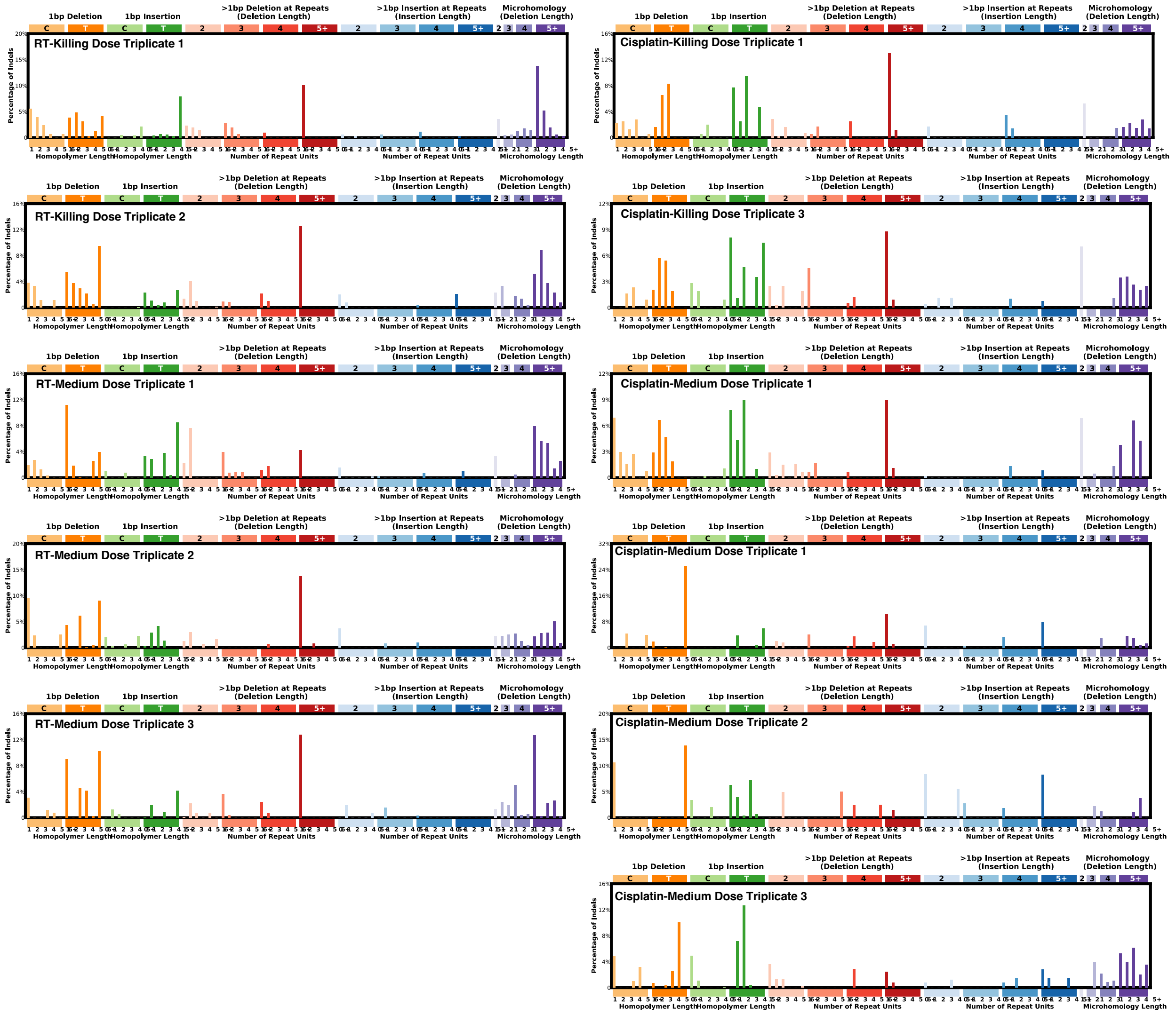

**Figure S7:** Median profiles of sublethal-dose platinum and RT single-cell-derived colonies.

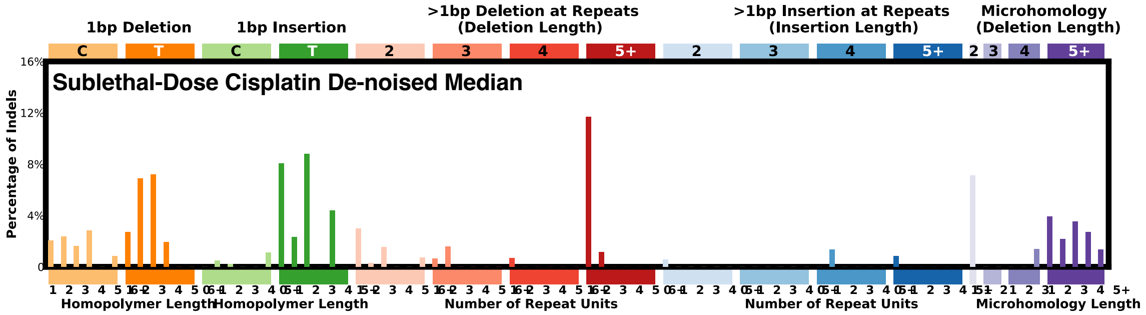


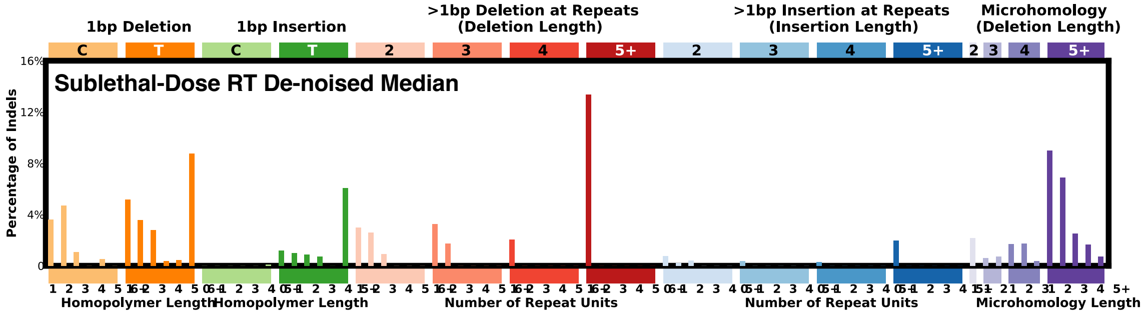

**Figure S8:** Absolute contribution of ID83 signatures from *mmsig* re-fitting of *de novo* AML and t-MN.

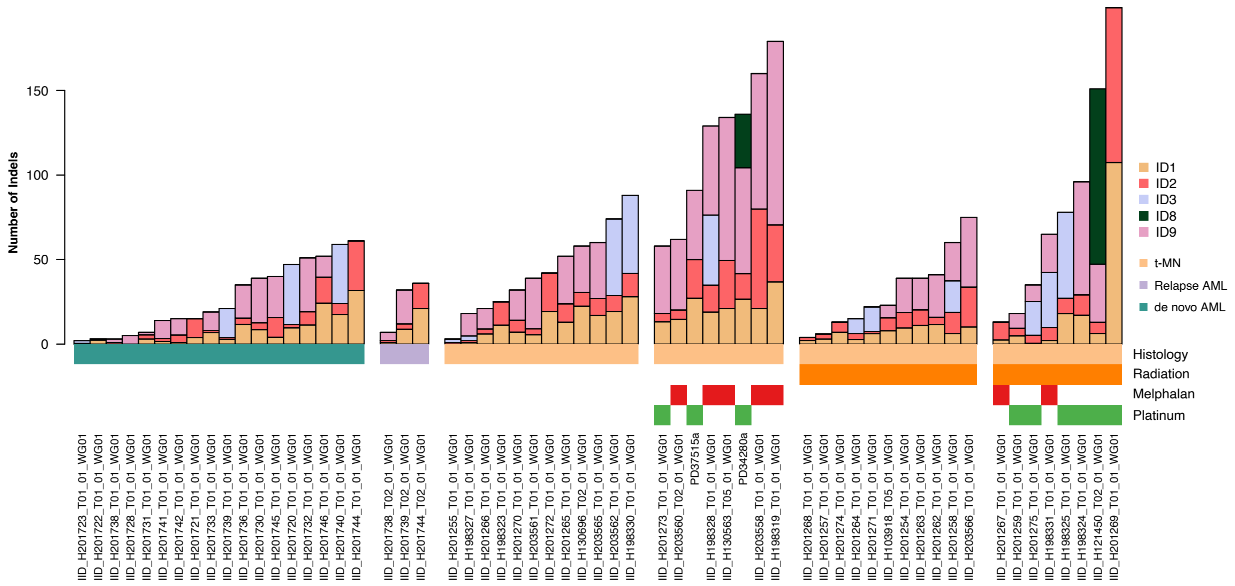
